# Spatially heterogeneous microstructural development within subcortical regions from 9-13 years

**DOI:** 10.1101/2021.06.04.446984

**Authors:** C E Palmer, D Pecheva, J Iversen, DJ Hagler, L Sugrue, P Nedelec, C Fan, W K Thompson, T L Jernigan, A M Dale

## Abstract

During late childhood behavioral changes, such as increased risk-taking and emotional reactivity, have been associated with the maturation of cortico-subcortical circuits. Understanding microstructural changes in subcortical regions may aid our understanding of how individual differences in these behaviors emerge. Restriction spectrum imaging (RSI) is a framework for modelling diffusion-weighted imaging that decomposes the diffusion signal from a voxel into hindered and restricted compartments. This yields greater specificity than conventional methods of characterizing intracellular diffusion. Using RSI, we modelled voxelwise restricted isotropic, N0, and anisotropic, ND, diffusion across the brain and measured cross-sectional and longitudinal age associations in a large sample (n=8,039) from the Adolescent Brain and Cognitive Development (ABCD) study aged 9-13 years. Older participants had higher N0 and ND across subcortical regions. The largest associations for N0 were within the basal ganglia and for ND within the ventral diencephalon. Importantly, age associations varied with respect to the internal cytoarchitecture within subcortical structures, for example age associations differed across thalamic nuclei. This suggests that developmental effects may map onto specific cell populations or circuits and highlights the utility of voxelwise compared to ROI-wise analyses. Future analyses will aim to understand the relevance of this subcortical microstructural developmental for behavioral outcomes.

## INTRODUCTION

Brain development during childhood and adolescence is associated with distributed structural alterations in both gray and white matter that occur concurrently with cognitive and behavioral development. Alterations in reward and affective processing are particularly pertinent during adolescence and associated with increased emotional reactivity (Casey et al., 2008). These behavioral changes are hypothesized to be underpinned by cortico-subcortical circuitry (Casey et al., 2016). The precise quantification of the microstructural changes during typical development may provide important information for understanding individual differences in cognition and the emergence of increased emotional reactivity and risk-taking in this period. Diffusion tensor imaging (DTI) has frequently been used to probe microstructural changes in the brain. Previous studies have shown increases in fractional anisotropy (FA) and decreases in mean diffusivity (MD) throughout the brain across childhood and into young adulthood, with variability in the trajectory of microstructural development across different brain regions (for review see Lebel & Deoni, 2018). Many studies have measured developmental changes in DTI metrics within white matter (WM) (Krogsrud et al., 2016; Catherine Lebel & Beaulieu, 2011; Pohl et al., 2016), but fewer studies have explored DTI changes in deep gray matter structures, in part due to the inadequacies of DTI for studying complex cytoarchitecture and the lower signal-to-noise ratio (SNR) when estimating FA in particular (Farrell et al., 2007). Despite this, in one study, increases in FA from 5-30 years appeared to be larger in subcortical regions compared to the WM tracts (Lebel et al., 2008).

However, the diffusion tensor model only allows the expression of a single principal direction of diffusion and is unable to characterize mixtures of neurite orientations within a voxel. Recent advances in diffusion data acquisition include multiple b-value acquisitions and high angular resolution diffusion imaging (HARDI) have enabled more complex models of tissue microstructure, taking into account multiple tissue compartments, multiple fiber populations in white matter and orientated structure of neurites within grey matter. Restriction spectrum imaging (RSI; Brunsing et al., 2017; White et al., 2013, 2014) uses multiple b-value HARDI data to model the diffusion-weighted signal as emanating from multiple tissue compartments, reflecting free, hindered and restricted water, with different intrinsic diffusion properties. The hindered compartment primarily represents extracellular space although may also describe diffusion within intracellular spaces with dimensions larger than the diffusion length scale (typically, ~10μm, for the diffusion sequences used in human imaging studies). The apparent diffusivity of the hindered compartment decreases with increasing tortuosity of the space, which is in turn related to the extracellular volume fraction and cell packing density (White et al., 2014). The restricted compartment primarily represents intracellular space, within cells or processes of dimensions smaller than the diffusion length scale. Free water diffusion primarily represents CSF or intravascular spaces. Within each voxel, RSI models the diffusion signal as a linear mixture of these different compartments. Spherical deconvolution (SD) is used to reconstruct the fiber orientation distribution (FOD) in each voxel for each compartment. There have been few developmental studies that have primarily focused on how restricted (intracellular) diffusion changes with age, therefore in the current study we have used RSI to specifically probe changes in restricted diffusion in late childhood. The spherical harmonic (SH) coefficients are used to estimate the relative signal fraction of restricted anisotropic (directional) diffusion (ND) and restricted isotropic (zeroth order) diffusion (N0). Unlike the standard tensor model, RSI can dissociate diffusion emanating from different oriented structures within the same voxel, for example crossing fibers. N0 can be modulated by both structures with a spherical or compact shape less than the diffusion length scale, such as neuronal cell bodies or microglia, within which diffusion is isotropic, and by multiple cylindrical structures in the same voxel oriented such that anisotropic diffusion is occurring in all directions. To use a signal processing analogy, ND is akin to the weighted sum of different components in the Fourier series, whereas N0 is analogous to the direct-current offset. Using this model, we can more precisely understand the underlying microstructural changes occurring during late childhood and adolescent development.

RSI has not previously been used to study developmental changes in late childhood. However, similar multi-compartment models, such as neurite orientation dispersion and density imaging (NODDI), have been shown to be more sensitive to developmental changes than DTI metrics (Genc et al., 2017). Across children aged 8-13 years, Mah et al. (2017) showed significant increases in neurite density index (NDI) from the NODDI model across the globus pallidus, putamen, thalamus, hippocampus and amygdala, but no changes in orientation dispersion index (ODI). NDI estimates the intracellular volume fraction, irrespective of orientation. ODI measures the degree of dispersion of intracellular diffusion. Therefore, these results suggest that developmental changes in the deep gray matter are associated with increases in myelination or the size/number of neurites rather than neurite coherence or geometry. Although NODDI is a useful model for describing intracellular diffusion, NDI is limited in that it represents a measure of the total intracellular volume fraction; in contrast, RSI can delineate isotropic and anisotropic diffusion within the restricted compartment, and can therefore provide different, more specific information about the development of intracellular diffusion within each voxel.

In the current study, we have used longitudinal data across two time points to estimate developmental changes in tissue microstructure within and surrounding subcortical regions. We have modeled both baseline age (age_0_) and change in age from baseline to follow-up (age_Δ_) in order to delineate cross-sectional and longitudinal developmental effects (Morrell et al., 2009; Sørensen et al., 2021). The cross-sectional effects estimate variability in our diffusion metrics with age across all individuals at a given point in time; whereas, the longitudinal effects estimate how the brain changes with increasing age within subjects with multiple time points. We have used data from Release 3.0 of the Adolescent Brain and Cognitive Development (ABCD) Study to measure voxelwise developmental changes in isotropic and anisotropic restricted (intracellular) diffusion. The large sample size (n=8,039) and small age range at each time point (9-11 years at baseline; 11-13 years at follow-up) provides high precision to delineate microstructural changes in subcortical gray matter regions and surrounding white matter.

## METHODS

### Sample

The ABCD study is a longitudinal study across 21 data acquisition sites following 11,880 children starting at 9-11 years. This paper uses baseline and two-year follow up (FU) data from the NIMH Data Archive ABCD Collection Release 3.0 (DOI: 10.15154/1519007). The ABCD cohort is epidemiologically informed (Garavan et al., 2018), including participants from demographically diverse backgrounds, and has an embedded twin cohort and many siblings. Exclusion criteria for participation in the ABCD Study were limited to: 1) lack of English proficiency in the child; 2) the presence of severe sensory, neurological, medical or intellectual limitations that would inhibit the child’s ability to comply with the protocol; 3) an inability to complete an MRI scan at baseline. The study protocols are approved by the University of California, San Diego Institutional Review Board. Parent/caregiver permission and child assent were obtained from each participant.

All statistical analyses included 12,525 observations with 8,039 unique subjects, such that 4,486 subjects had data at two time points. Observations were included in the final sample if the participant had complete data across sociodemographic factors (household income, highest parental education, ethnicity), available genetic data (to provide ancestry information using the top 10 principal components), available imaging data that passed all diffusion inclusion criteria and available information regarding acquisition scanner ID and software version. In the ABCD Study, Release 3.0, there are 16727 available scans with scanner information (9.3% missingness). Of these scans, 1728 were excluded for not meeting the recommended dMRI inclusion criteria outlined in the Release 3.0 release notes (tabulated variable: imgincl_dmri_include = 0) and an additional 20 observations were excluded for poor registration defined below (in *Atlas Registration*). The final sample included all remaining observations that had complete data for the previously listed information. Table 1 shows the demographics of the final sample used for statistical analysis stratified by event. Participants who had competed their 2 year follow up in Release 3.0 were more likely to have higher household income, higher parental education and have male assigned as their sex at birth. This may reflect differences in recruitment procedures over the course of recruitment in order to ensure the final sample reflected the demographics of the US population as closely as possible.

**Table 1.**
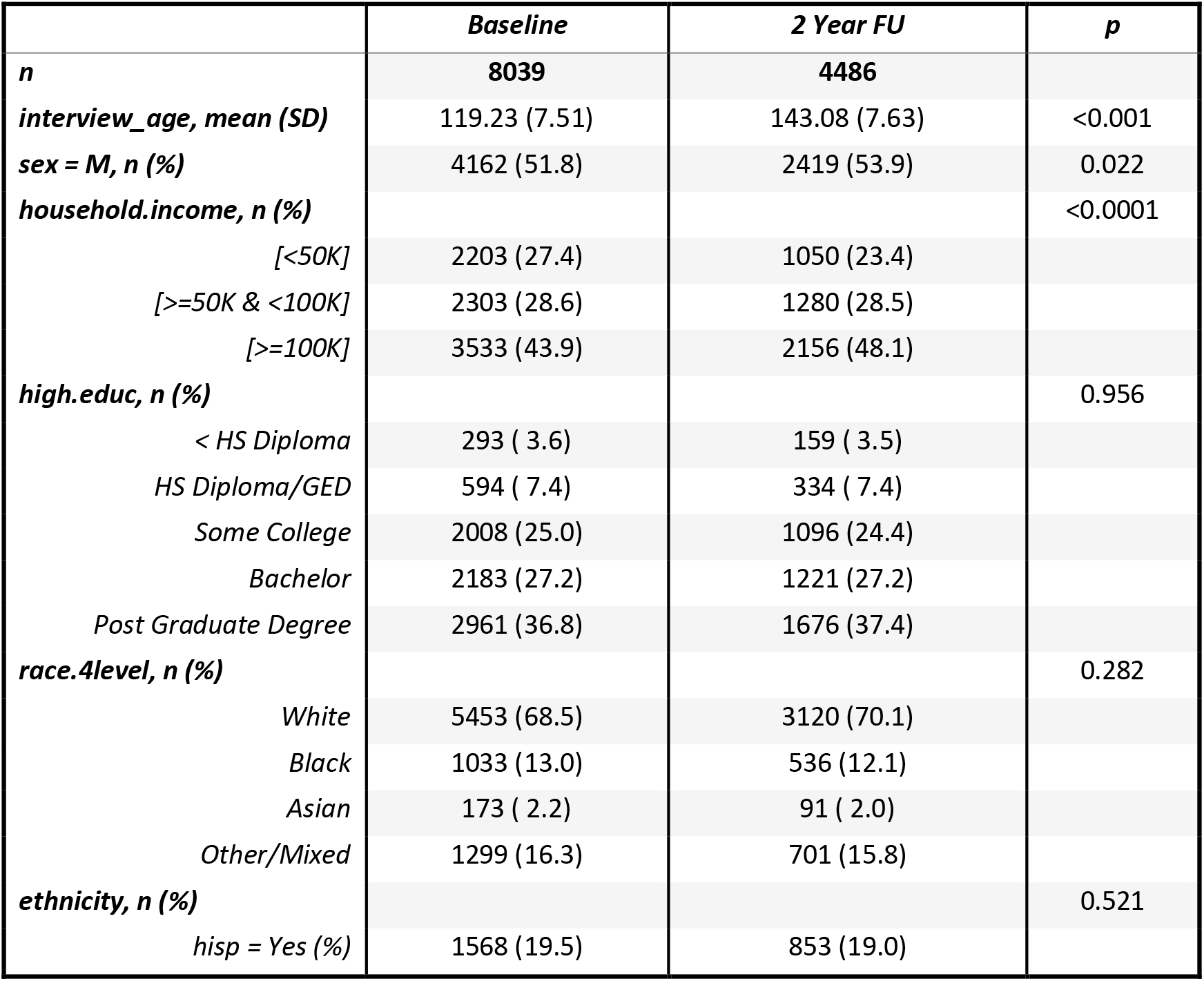
Demographics of the sample. Demographic data is shown for age (mean, (SD)), sex at birth, household income, parental education, self-declared race and endorsement of Hispanic ethnicity (n, (%)). These factors are stratified by time point: baseline and 2-year FU. There were significant differences in the socioeconomic status (income and parental education) and sex at birth for those who had 2-year follow up data in Release 3.0 indicative of differences in the demographics of participants as they were recruited. Participants recruited earlier in the study were more likely to have higher household income and parental education and be born male. All of these variables are controlled for in all statistical analyses to account for this.

|  | *Baseline* | *2 Year FU* | *p* |
| --- | --- | --- | --- |
| *n* | **8039** | **4486** |  |
| *interview_age, mean (SD)* | 119.23 (7.51) | 143.08 (7.63) | <0.001 |
| *sex = M, n (%)* | 4162 (51.8) | 2419 (53.9) | 0.022 |
| *household.income, n (%)* |  |  | <0.0001 |
| [<50K] | 2203 (27.4) | 1050 (23.4) |  |
| [>=50K & <100K] | 2303 (28.6) | 1280 (28.5) |  |
| [>=100K] | 3533 (43.9) | 2156 (48.1) |  |
| *high.educ, n (%)* |  |  | 0.956 |
| < HS Diploma | 293 ( 3.6) | 159 ( 3.5) |  |
| HS Diploma/GED | 594 ( 7.4) | 334 ( 7.4) |  |
| Some College | 2008 (25.0) | 1096 (24.4) |  |
| Bachelor | 2183 (27.2) | 1221 (27.2) |  |
| Post Graduate Degree | 2961 (36.8) | 1676 (37.4) |  |
| *race.4level, n (%)* |  |  | 0.282 |
| White | 5453 (68.5) | 3120 (70.1) |  |
| Black | 1033 (13.0) | 536 (12.1) |  |
| Asian | 173 ( 2.2) | 91 ( 2.0) |  |
| Other/Mixed | 1299 (16.3) | 701 (15.8) |  |
| *ethnicity, n (%)* |  |  | 0.521 |
| hisp = Yes (%) | 1568 (19.5) | 853 (19.0) |  |

### MRI acquisition

The ABCD MRI data were collected across 21 research sites using Siemens Prisma, GE 750 and Philips 3T scanners. Scanning protocols were harmonized across sites. Full details of structural and diffusion imaging acquisition protocols used in the ABCD study have been described previously (Casey et al., 2018; Hagler et al., 2019) so only a short overview is given here. dMRI data were acquired in the axial plane at 1.7mm isotropic resolution with multiband acceleration factor 3. Diffusion-weighted images were collected with seven b=0 s/mm^2^ frames and 96 non-collinear gradient directions, with 6 directions at b=500 s/mm^2^, 15 directions at b=1000 s/mm^2^, 15 directions at b=2000 s/mm^2^, and 60 directions at b=3000 s/mm^2^. T1-weighted images were acquired using a 3D magnetization-prepared rapid acquisition gradient echo (MPRAGE) scan with 1mm isotropic resolution and no multiband acceleration. 3D T2-weighted fast spin echo with variable flip angle scans were acquired at 1mm isotropic resolution with no multiband acceleration.

### Image Processing

The processing steps for diffusion and structural MR data are outlined in detail in Hagler et al., (2019). Briefly, dMRI data were corrected for eddy current distortion using a diffusion gradient model-based approach (Zhuang et al., 2006). To correct images for head motion, we rigid-body-registered each frame to the corresponding volume synthesized from a robust tensor fit, accounting for image contrast variation between frames. Dark slices caused by abrupt head motion were replaced with values synthesized from the robust tensor fit, and the diffusion gradient matrix was adjusted for head rotation (Hagler et al., 2009, 2019). Spatial and intensity distortions caused by B0 field inhomogeneity were corrected using FSL’s *topup* (Andersson et al., 2003) and gradient nonlinearity distortions were corrected for each frame (Jovicich et al., 2006). The dMRI data were registered to T1w structural images using mutual information (Wells et al., 1996) after coarse pre-alignment via within-modality registration to atlas brains. dMRI data were then resampled to 1.7 mm isotropic resolution, equal to the dMRI acquisition resolution.

T1w and T2w structural images were corrected for gradient nonlinearity distortions using scanner-specific, nonlinear transformations provided by MRI scanner manufacturers (Jovicich et al., 2006; Wald et al., 2001) and T2w images are registered to T1w images using mutual information (Wells et al., 1996). Intensity inhomogeneity correction was performed by applying smoothly varying, estimated B1-bias field (Hagler et al., 2019). Images were rigidly registered and resampled into alignment with a pre-existing, in-house, averaged, reference brain with 1.0 mm isotropic resolution (Hagler et al., 2019).

### Microstructural models

#### Restriction spectrum imaging (RSI)

The RSI model was fit to the diffusion data to model the diffusion properties of the cerebral tissue (White et al., 2013, 2014). RSI estimates the relative fraction that separable pools of water within a tissue contribute to the diffusion signal, based on their intrinsic diffusion characteristics. Free water (e.g., CSF) is defined by unimpeded water diffusion. Hindered diffusion follows a Gaussian displacement pattern characterised by the presence of neurites (axons and dendrites), glia and other cells. This includes water both within the extracellular matrix and certain intracellular spaces with dimensions larger than the diffusion length scale (typically, ~10μm, for the diffusion sequences used in human imaging studies (White et al., 2013)). Restricted diffusion follows a non-Gaussian pattern of displacement and describes water within intracellular spaces confined by cell membranes. Imaging scan parameters determine the sensitivity of the diffusion signal to diffusion within these separable pools. At intermediate b-values (b=500-2500s/mm^2^), the signal is sensitive to both hindered and restricted diffusion; whereas, at high b-values (b=3000s/mm^2^), the signal is primarily sensitive to restricted diffusion. The hindered and restricted compartments are modeled as fourth order spherical harmonic (SH) functions and the free water compartment is modelled using zeroth order SH functions. The longitudinal diffusivity (LD) is held constant, with a value of 1 × 10^−3^ mm^2^/s for the restricted and hindered compartments. For the restricted compartment, the transverse diffusivity (TD) is fixed to 0 mm^2^/s. For the hindered compartment, TD is fixed to 0.9 × 10^−3^ mm^2^/s. For the free water compartment the isotropic diffusivity is fixed to 3 × 10^−3^ mm^2^/s. Theoretically, any increases in the tortuosity of the hindered compartment, for example due to a decrease in the volume of the extracellular space, will decrease the effective diffusivity in the hindered compartment; however, in our model we are assuming the hindered diffusivity is constant. Spherical deconvolution (SD) is used to reconstruct the fiber orientation distribution (FOD) in each voxel from the restricted compartment. The restricted anisotropic measure, ND, is the norm of the SH coefficients for the second and fourth order SHs (divided by the norm of all the coefficients across the restricted, hindered and free water compartments). This models oriented diffusion emanating from multiple directions within a voxel. The restricted isotropic measure, N0, refers to the spherical mean of the FOD across all orientations (zeroth order SH divided by the norm of all the coefficients across the restricted, hindered and free water compartments).

In this study we explore associations between age and the rotation-invariant features of the restricted compartment FOD. For a detailed description of the derivation of the RSI model see White et al. (2013). We extracted a measure of the restricted isotropic and restricted anisotropic diffusion signal. Within each voxel the total diffusion signal, S, can be represented as

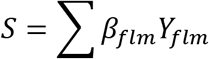

where *Y*_*flm*_ is a SH basis function of order *l* and degree *m* of the FOD corresponding to the *f* th compartment, and *β*_*flm*_ are the corresponding SH coefficients. The measure of restricted isotropic diffusion is given by the coefficient of the zeroth order SH coefficient, *β*_*f*,*l*=0,*m*=0_, normalized by the Euclidian norm of all *β*_*flm*_ and termed N0:

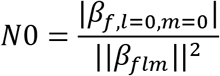

The measure of restricted normalized anisotropic diffusion is given by the norm of *β*_*flm*_, where *l* > 0, and *f* is the restricted compartment, and is termed ND:

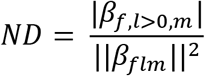

These normalized RSI measures are unitless and range from 0 to 1. As N0 and ND are normalized with respect to the hindered and free water compartments, changes in restricted diffusion will be relative to the other compartments. The relative size and shape of restricted compartments will differentially modulate isotropic and anisotropic diffusion.

The magnitude of diffusion that we are sensitive to is dependent on the diffusion scan parameters. Typical diffusion times used in clinical DWI scans are approximately 10–50ms corresponding to average molecular displacements on the order of 10μm (Mukherjee et al., 2008). Any water displacements smaller than this scale would not lead to changes in the measured diffusion coefficient, but changes in cell size <~10μm could alter the relative signal fractions of hindered and restricted diffusion in a voxel. Diffusion estimated in these compartments is also dependent on the permeability of cellular membranes; greater exchange across intracellular and extracellular space will mean that diffusion will appear more hindered rather than restricted. Supplementary Table 1 outlines the expected changes to hindered and restricted diffusion following example microstructural developmental processes.

#### Diffusion tensor imaging

The diffusion tensor model (Basser et al., 1994; Basser & Pierpaoli, 1996) was used to calculate fractional anisotropy (FA) and mean diffusivity (MD). Diffusion tensor parameters were calculated using a standard, linear estimation approach with log-transformed diffusion-weighted (DW) signals (Basser et al., 1994). Tensor matrices were diagonalized using singular value decomposition, obtaining three eigenvectors and three corresponding eigenvalues. FA and MD were calculated from the eigenvalues (Basser & Pierpaoli, 1996).

### Atlas registration

To allow for voxelwise analysis, subjects’ imaging data were aligned using a multimodal nonlinear elastic registration algorithm. At the end of the preprocessing steps outlined in *Image Processing* and described in detail in Hagler et al. (2019), subjects’ structural images and diffusion parameter maps were aligned to an ABCD-specific atlas, using a custom diffeomorphic registration method. The ABCD-specific atlas was constructed using an iterative procedure, consisting of an initial affine registration, followed by a multi-scale, multi-channel elastic diffeomorphic registration. Imaging volumes used for the elastic registration included 3D T1 and direction-encoded color (DEC) FA volumes (x-, y-, and z-orientations). After each iteration, morphed volumes for each subject were averaged to create an updated atlas, and then the process was repeated until convergence. Participants with poor registration to atlas were excluded from the average and subsequent statistical analyses. Poor registration was defined as a voxelwise correlation to atlas <0.8 (see *Sample* for number excluded).

### Labelling regions of interest (ROI)

Subcortical structures were labeled using Freesurfer 5.3 (Fischl et al., 2002). Subjects’ native space Freesurfer parcellations were warped to the atlas space and averaged across subjects. Bilateral binary masks for each ROI were created using a probabilistic threshold of 0.8. Additional subcortical nuclei, not available in the FreeSurfer segmentation, were labeled by registering readily available, downloadable, high spatial resolution atlases to our atlas space. The Pauli atlas was generated using T1 and T2 scans from 168 typical adults from the Human Connectome Project (HCP) (Pauli et al., 2018). The Najdenovska thalamic nuclei atlas was generated using a k-means algorithm taking as inputs the mean FOD SH coefficients from within a Freesurfer parcellation of the thalamus, using adult HCP data from 70 subjects (Najdenovska et al., 2018). Bilateral binary masks were created for all ROIs in atlas space. All subcortical ROIs and abbreviations are listed in Supplementary Table 2.

### Statistical analysis

Univariate general linear mixed effects models (GLMMs) were applied at each voxel to test the associations between age and diffusion metrics: N0, ND, FA and MD. Age was divided into two predictors: baseline age (age_0_); and, change in age from baseline to 2 year follow up (age_Δ_). A contrast (age_0_=1, age_Δ_=-1) was used to measure significant differences in the voxelwise estimated beta coefficients for age_0_ and age_Δ_. Interactions between each age predictor and sex were included in the model: age_0_*sex; age_Δ_*sex. Given the demographic diversity in the sample, all statistical analyses controlled for the variables shown in Table 1, except fixed effects of the top 10 genetic principal components were used to account for ancestry effects in lieu of self-declared race. Additional fixed effects included scanner ID and MRI software version. Random effects of family ID and subject ID were also modelled. Whole-brain voxelwise analyses were corrected for multiple comparisons at an alpha level of 0.05 using a Bonferroni correction across 155,179 voxels (threshold for voxelwise significance: |t|=4.98, uncorrected −log_10_(p)=6.49). This provides a conservative estimate of significant developmental effects as the true number of independent tests is likely to be smaller than this. T-statistic maps thresholded to correct for multiple comparisons across the whole brain are presented in the supplementary materials. Unthresholded effect size maps are shown in the main figures to provide a comprehensive description of the distribution of estimated effects across the brain. ROI analyses were also conducted using the same GLMMs by taking the mean diffusion metric across the voxels within each ROI mask as the dependent variable in order to provide an estimate of the developmental effect for each subcortical region. Violin plots were generated to show the variability in voxelwise effects across all voxels within each ROI mask in order to highlight the heterogeneity of developmental effects within each ROI.lp;. P-values were adjusted for multiple comparisons at an alpha level of 0.05 based on 22 independent tests (threshold for ROIwise significance: |t|=2.83, uncorrected - log_10_(p)=2.64, corrected −log_10_(p)=1.3).

## RESULTS

### Developmental effects of restricted isotropic diffusion, N0, across subcortical regions

N0 was positively associated with age across the brain with the greatest effects in subcortical regions, particularly within the basal ganglia (age_0_: max *β*=0.013, t=25, −log_10_(p-adj)=135; age_Δ_: max *β* =0.012, t=34, −log_10_(p-adj)=255; see supplementary table 2 for summary statistics in each ROI). The patterns of association across subcortical regions for age_0_ and age_Δ_ were very similar (Figure 1; column 1-2), but the t-statistics were much greater for ageΔ due to the increased statistical power when estimating longitudinal effects. Voxels showing statistically significant associations corrected for multiple comparisons are shown in Supplementary Figure 1A-D. There were no significant voxels showing a significant difference between the beta coefficients for age_0_ and age_Δ_ after correction for multiple comparisons. There were no significant voxelwise age_0_ or age_Δ_ by sex interactions for N0. Voxelwise mean N0, averaged across participants, varied across subcortical gray and WM (Figure 1; column 3); however, larger age associations were not only found in regions of high mean N0. Voxelwise FODs, averaged across participants, show the orientation structure of diffusion in each voxel and are colored based on the diffusion direction (Figure 1; column 4). There was clear variability in the orientation structure of diffusion within gross ROIs and the surrounding WM. By including atlases with a finer subcortical parcellation, we were able to localize developmental effects within large subcortical structures.

**Figure 1.**
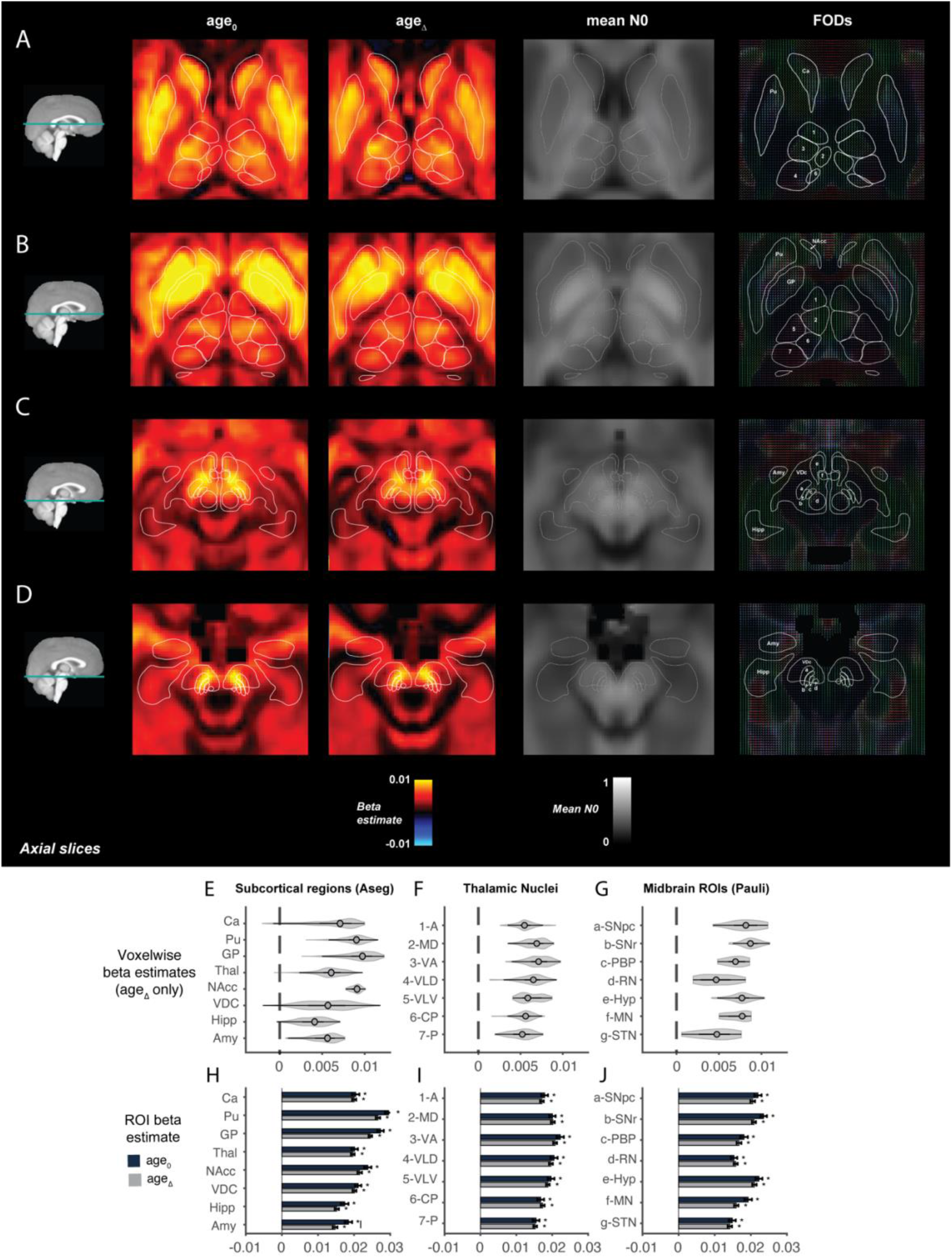
Associations between age_0_ and age_Δ_ and restricted isotropic diffusion, N0, across subcortical regions. A-C) Voxelwise beta coefficients for the association between N0 and age_0_ and age_Δ_ across different axial brain slices moving from superior (top) to inferior (bottom; columns 1 and 2); N0 averaged across all subjects in the same brain slices (column 3); voxelwise FODs averaged across all subjects in the same brain slices (column 4). Outlines of the Aseg, Pauli and Najdenovska ROIs are overlaid. E-G) Violin plots showing the distribution of voxelwise age_Δ_ associations in each ROI. H-J) ROI analyses of mean N0 in each ROI associated with age_0_ (dark blue) and age_Δ_ (gray). Asterisks represent significant associations (corrected p-value threshold of −log_10_(p)=2.64). Solid lines represent significant difference between age_0_ and age_Δ_ coefficients.

Voxelwise age effects were heterogeneous in magnitude across and within subcortical regions. Voxels in the globus pallidus (GP) and putamen, the surrounding WM between and ventral to these structures, and voxels within the ventral diencephalon (VDC) showed the largest age associations (Figure 1A-D). Voxelwise associations were the most heterogeneous across the VDC (Figure 1E). Within the VDC, voxels showing the largest N0 associations were within the posterior portions of the hypothalamus, the substantia nigra pars compacta (SNpc), substantia nigra pars reticulata (SNpr) and WM between those regions (Figure 1G). These associations did not necessarily coincide with regions of highest mean N0. Age associations were found across the thalamic nuclei (Figure 1F) with the largest associations in more anterior nuclei particularly along the lateral edge of the ventral anterior nucleus (VA) coinciding with higher mean N0. ROI analyses, reflecting age associations with mean N0 within each region, showed similar positive effects for age_0_ and age_Δ_ across all subcortical ROIs (Figure 1H-J). Across ROIs, the amygdala was the only region in which the age_Δ_ beta estimate was significantly less than the age_0_ beta estimate, suggesting a divergence from linearity from 9-13 years in this region. The putamen, GP, nucleus accumbens and hippocampus also showed a nominal reduction in the age_Δ_ effect, however the difference in the beta coefficient did not reach the corrected threshold for significance (supplementary table 2).

Figure 3 shows images of the largest age_Δ_ associations zoomed in on specific ROIs in order to highlight examples of how these associations map onto changes in diffusion orientation. Figure 3A shows a coronal view of an area of N0 age_Δ_ associations extending ventral to the GP overlaying with diffusion occurring primarily in the lateral-medial (L-M) direction, which is likely to represent the anterior commissure; however this location is difficult to distinguish from the ventral pallidum (VP), which sits below the anterior commissure. When looking in the sagittal view (figure 3B), we can see that these associations extend through the extended amygdala, nucleus accumbens and head of the caudate. These effects appear to map onto voxels with diffusion in both the L-M and anterior-posterior (A-P) direction. The largest N0 development effects in the VDC were seen in voxels with diffusion primarily in the A-P direction (Figure 3C-D). Across the thalamus, effects were larger in anterior nuclei where diffusion was also primarily in the A-P direction and along the lateral edge of the VA nucleus where there was increased mean isotropy and diffusion crossing in multiple directions (Figure 3E).

### Developmental effects of restricted isotropic diffusion, ND, across subcortical regions

Across the brain, greater ND was associated with older age. The largest effects were found within the VDC (age_0_: max *β*=0.009, t=15, −log_10_(p-adj)=46; age_Δ_: max *β* =0.009, t=23, −log_10_(p-adj)=113; see supplementary table 3 for summary statistics in each ROI). As with N0, the maps of beta coefficients were very similar for age_0_ and age_Δ_ (figure 2; column 1-2), but the t-statistics were much greater for age_Δ_ due to the increased statistical power when estimating longitudinal effects. Voxels showing statistically significant associations, corrected for multiple comparisons, are shown in Supplementary Figure 1E-H. No voxels survived correction for multiple comparison across the brain for the contrast between age_0_ and age_Δ_. There were no significant voxelwise age_0_ or age_Δ_ by sex interactions on ND. Voxelwise mean ND, averaged across participants, varied across subcortical gray and WM with the highest values in the WM (figure 2; column 3); again, larger age associations were not only found in regions of high mean ND. Voxelwise FODs, averaged across participants, show the orientation structure of diffusion in each voxel and are colored based on the diffusion direction (Figure 2; column 4).

**Figure 2.**
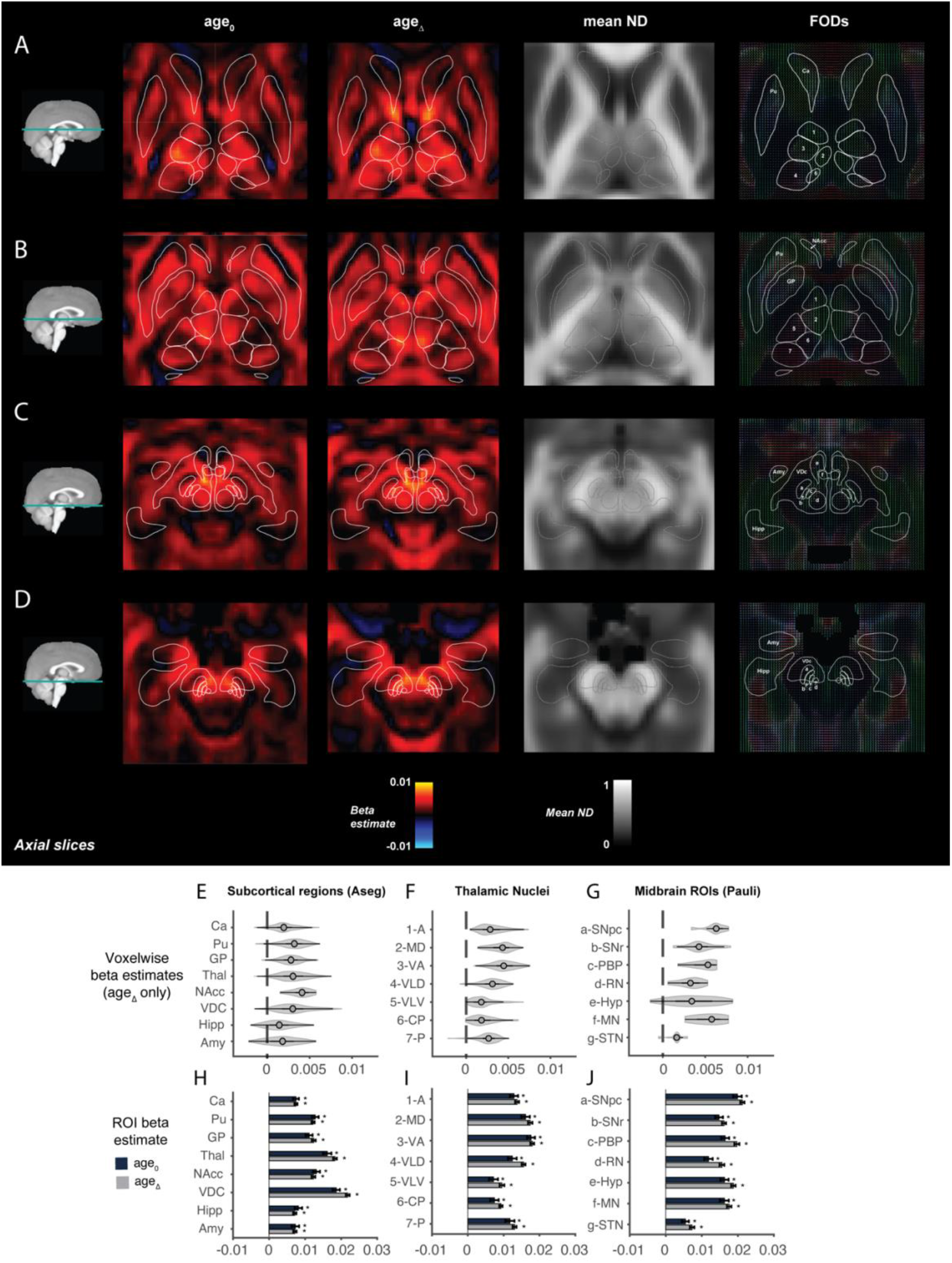
Associations between age_0_ and age_Δ_ and restricted anisotropic diffusion, ND, across subcortical regions. A-C) Voxelwise beta coefficients for the association between ND and age_0_ and age_Δ_ across different axial brain slices moving from superior (top) to inferior (bottom; columns 1 and 2); ND averaged across all subjects in the same brain slices (column 3); voxelwise FODs averaged across all subjects in the same brain slices (column 4). Outlines of the Aseg, Pauli and Najdenovska ROIs are overlaid. E-G) Violin plots showing the distribution of voxelwise age_Δ_ associations in each ROI. H-J) ROI analyses of mean N0 in each ROI associated with age_0_ (dark blue) and age_Δ_ (gray). Asterisks represent significant associations (corrected p-value threshold of −log_10_(p)=2.64). Solid lines represent significant difference between age_0_ and age_Δ_ coefficients.

Developmental effects on ND were smaller in magnitude and more heterogeneous across voxels within subcortical regions than for N0. Within the putamen there was a lack of effects in voxels around the medial edge. Within the caudate positive age associations were only found in the head of the caudate along the ventral edge closest to the thalamus extending posteriorly and ventrally outside of the caudate ROI boundary into the WM anterior to the thalamus and towards the ventral striatum (Figure 2A-B; Figure 3B). Effects were smaller in the putamen and caudate where mean ND across subjects was low. Mean ND was also low within the amygdala and hippocampus; positive age associations were only found within medial anterior portions (Figure 2C-D). The largest age associations were primarily in voxels in the midbrain region, including such nuclei as the SN and a posterior portion of the hypothalamus (Figure 2C-D; Figure 3C-D).

**Figure 3.**
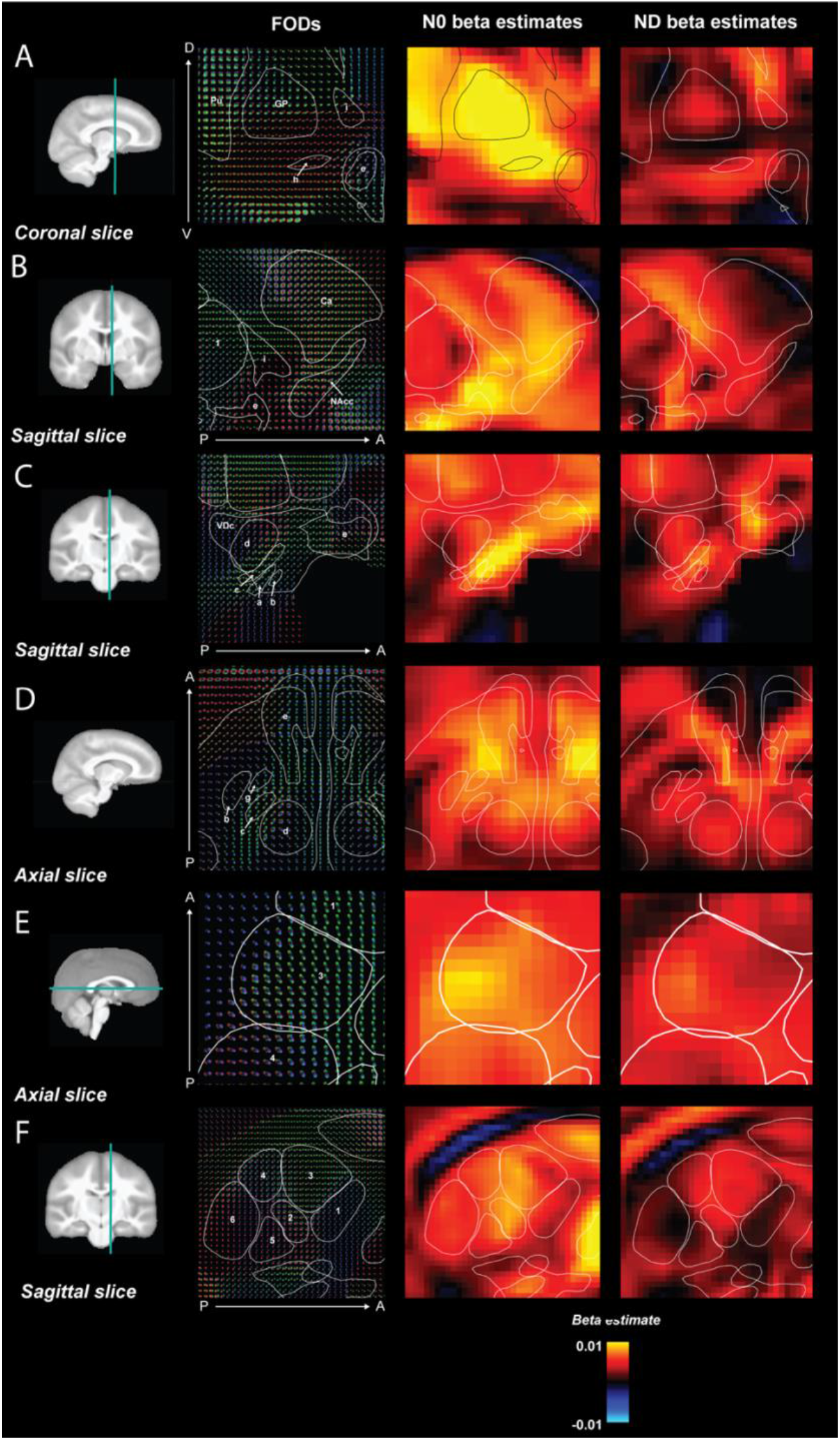
Zoomed in images of the voxelwise age_Δ_ associations with N0 and ND and FODs across different brain slices. A) Coronal view of the GP, VP(h), EA(i), and Hyp showing large N0 effects extending from the GP into the ventral region encompassing the ventral pallidum and anterior commissure with primarily lateral-medial (L-M; red) diffusion; B) Sagittal view showing associations within the Ca extending into the ventral striatum, EA and Hyp with both L-M and anterior-posterior (A-P; green) diffusion; C) Sagittal view of the VDC with ROIs of the RN (d), PBP (c), SNpc (a), SNpr (b) and Hyp (e) overlaid. The largest N0 associations here can be seen in the SNpc and SNpr extending into the surrounding WM along voxels with diffusion primarily in the A-P and dorsal-ventral (D-V, blue) direction. D) Axial view of the ventral diencephalon showing, for both N0 and ND, larger age associations in the posterior Hyp. E) Axial view of the VA nucleus of the thalamus (3) showing both N0 and ND associations in voxels with crossing diffusion directions. F) Sagittal view of the thalamic nuclei showing the largest associations for N0 and ND in the VA nucleus (3) where diffusion is primarily in the anterior-posterior (A-P, green) direction. There were no significant associations between ND and age_Δ_ in voxels within the central and pulvinar nuclei where diffusion was primarily in the L-M direction.

There was variability in the voxelwise effects across all subcortical regions, with the greatest heterogeneity in the VDC, thalamus and hypothalamus (Figure 2E-G). Within the thalamus, there were a number of voxels within the anterior, ventrolateral and pulvinar nuclei that showed no significant age association. The largest positive age associations were found in the VA and medial dorsal (tMD) nuclei (Figure 2F). The heterogeneity in age effects across the thalamus was particularly clear when looking at the ROI analysis for each nucleus (Figure 2I). Across the thalamus and VDC, the largest effects seemed to map onto voxels with diffusion occurring primarily within the anterior-posterior direction (Figure 3C-F). ROI analyses showed similar beta estimates for age_0_ and age_Δ_ across all subcortical ROIs (Figure 2H-J; supplementary table 3).

### Developmental associations with DTI vs RSI metrics

Supplementary figure 3 shows the voxelwise and ROI-wise age_0_ and age_Δ_ associations with mean diffusivity (MD) and fractional anisotropy (FA) from the diffusion tensor model. Supplementary figure 4 shows the thresholded voxelwise t-statistics. In general, there was a strong but inverse correspondence between the MD and N0 developmental associations across the white matter and subcortical ROIs; predominantly in regions where N0 was positively associated with age, MD was negatively associated with age. However, there were subtle differences in the magnitude of effects across ROIs highlighting the different models used to estimate these measures. There were larger differences between the FA and ND associations. Namely, the magnitude of the FA associations was much smaller than ND, such that a larger sample would be required to detect developmental FA associations. This was particularly true for the age_0_ associations in the thalamus where very few voxels showed significant FA age associations.

## DISCUSSION

In this study, we have shown statistically significant age associated increases in restricted (primarily intracellular) diffusion across subcortical gray and WM in a large sample (n=8,039) of children from 9 to 13 years. This is the largest study to date measuring both cross-sectional and longitudinal changes in diffusion metrics at this age and utilizing novel RSI measures. By modelling age_0_ and age_Δ_ as independent predictors we were able to delineate both between and within-subject structural associations with age. In this age range, these effects were very similar, suggesting a generally linear relationship between age and restricted water diffusion in subcortical regions. Across the brain, the largest age-related increases in isotropic, N0, and anisotropic, ND, diffusion were detected in the basal ganglia, VDC, and surrounding WM. Voxelwise age associations were highly variable within subcortical regions and the boundaries of these significant effects mapped onto changes in the orientation of the FODs. This suggests that we can identify dissociable developmental effects within subcomponents of subcortical structures that may be associated with differing functional circuits. This highlights the benefit of measuring voxelwise compared to ROI-wise associations and utilizing high resolution parcellations of subcortical structures that reflect known histologic/functional subdivisions within deep gray matter nuclei such as the thalamus.

Previous studies have highlighted significant changes in FA and MD across subcortical regions (Baron Nelson et al., 2019; Lebel et al., 2008; Simmonds et al., 2014), with regions of the basal ganglia showing greater percentage change from 5-30 years than many WM tracts (Lebel et al., 2008), in agreement with the results reported here. From 8-13 years, Mah et al (2017) found that NDI from the NODDI model, a measure of the intracellular volume fraction, showed the largest percent increase in the pallidum (10-13% change) followed by the putamen, hippocampus, amygdala and thalamus (3-7% change) and found no age association in the caudate. Although the RSI and NODDI models are very different, NDI, similar to N0, captures the total intracellular volume fraction in a voxel, therefore is most closely related to our measure of N0. As the intracellular volume fraction increases in a voxel, the magnitude of water displacement reduces, thereby decreasing MD. Indeed, NDI has previously been shown to correlate negatively with MD (Zhang et al., 2012), and in the current study MD showed age associations in the opposite direction to N0. Indeed, our N0 results were very similar to the NDI effects reported by Mah et al., apart from a significant age association in the caudate. This may reflect the greater sensitivity of the RSI model parameters to age-related changes in cytoarchitecture of the caudate and/or increased statistical power in this study to detect an association.

There are a number of microstructural changes associated with the neurobiological underpinnings of development that may contribute to the age-related increases in restricted diffusion detected in the current study. These are outlined in detail in Supplementary table 1. Subcortical gray matter has a complex cytoarchitecture consisting of many cell bodies of stellate shape, dendrites, terminal arbors and synapses that do not follow a layered structure. Developmental increases in the restricted signal fraction within the gray matter could be driven by increases in neurite density, dendritic sprouting or increases in cell size less than the typical diffusion length scale. At this scale, increasing cell size will not alter the magnitude of intracellular water displacement measurable, but will reduce the volume of the extracellular space. Given that restricted N0 and ND are normalized measures, changes in these metrics will always be relative to changes in the hindered and free water compartments, therefore any decreases in the hindered signal fraction will increase the restricted signal fraction increasing both N0 and ND. Indeed, in many voxels, N0 and ND both increased with age potentially reflecting neurobiological processes that reduce the hindered signal fraction. The relative size and shape of restricted compartments will then differentially modulate isotropic and anisotropic diffusion. For example, if a voxel has many neurites that are systematically aligned, an increase in the restricted signal fraction will have a larger impact on ND; however, if there are many dendrites in multiple directions, an increase in the restricted signal fraction will have a greater impact on N0. Areas in which N0 and ND dissociate provide interesting differential information about the specific microstructural changes occurring, such as in the main body of the caudate, which showed significant N0 associations, but no ND associations. The caudate has a very heterogeneous cytoarchitecture with many round islands of cells containing densely packed cells in variable size, shape and orientation, and a loosely packed matrix (Goldman-Rakic, 1982). The developmental isotropic, not anisotropic, microstructural changes in this region may reflect this cytoarchitecture.

By using voxelwise analyses we were able to measure the heterogeneity of developmental effects within subcortical regions highlighting the benefit of using voxelwise compared to ROI-wise analyses. We registered the Pauli and Najdenovska atlases to our data in order to utilize high-resolution finer subcortical parcellations to localize developmental changes within subcortical structures such as the thalamus. There was a clear pattern of developmental effects across the different thalamic nuclei, particularly for ND. Najdenovska et al (2018) generated the thalamic nuclei ROIs by clustering contiguous voxels with similar orientation microstructure determined by the FODs validating their results against a histological atlas (Najdenovska et al., 2018). When overlaying these ROIs on the average FODs measured in our sample, we could see (Figure 2) that the boundaries of the different nuclei indeed adhere to changes in the orientation of the average FODs. Within anterior nuclei (A, VA, tMD), where developmental effects were greatest for N0 and ND, diffusion primarily occurred in the anterior-posterior orientation (green), whereas within posterior nuclei (VLV, VLD, C, P), diffusion primarily occurred within the medial-lateral (red) orientation. The tMD nucleus of the thalamus is reciprocally interconnected with the prefrontal cortex and receives input from striatal, medial temporal, midbrain and basal forebrain structures (Groenewegen, 1988; Groenewegen et al., 1993; Ray & Price, 1993; Tanaka, 1976; Tobias, 1975; Vertes et al., 2015), therefore is well positioned to play a modulatory role within fronto-striatal-thalamo-cortical circuits thought to be important for several cognitive and emotional processes (Haber & Calzavara, 2009; Mitchell & Chakraborty, 2013; Ouhaz et al., 2018). Structural and functional connectivity of these thalamo-cortical connections has been shown to increase across childhood and adolescence (Alkonyi et al., 2011; Fair, 2010), and is thought to underpin behavioral changes in cognitive control and emotional reactivity during adolescence. Moreover, there were significant developmental effects on N0 in the region ventral to the GP and Ca, which encompasses both the ventral pallidum, ventral striatum (nucleus accumbens and olfactory tubercle) and bed of the nucleus stria terminalis (often referred to as the extended amygdala, as well as the anterior commissure (Zaborszky et al., 2015). These regions are highly interconnected with subcortical and cortical regions, particularly in frontal cortex, creating circuits integral for incentive-based learning, reward processing and decision-making (Barkley-Levenson & Galván, 2014; Delgado, 2007; Haber & Knutson, 2010). Microstructural changes within the thalamus and the ventral forebrain may be indicative of the refinement of these circuits in late childhood.

There were also statistically robust and heterogeneous associations within the VDC. The VDC is a group of structures that are poorly defined on T1w imaging, however, by calculating the mean voxelwise FODs across subjects, we could clearly see variability in the orientation structure of diffusion within this gross region highlighting the presence of potentially distinct cell populations. Changes in the orientation of the FODs also appeared to adhere to estimated outlines of finer subcortical parcellations that include many of the nuclei within the VDC from the Pauli atlas (Pauli et al., 2018). Indeed, the strongest associations between age and N0 and ND were in voxels oriented primarily in the anterior-posterior direction within and around the SN adjacent ventral tegmental area (VTA), which may reflect microstructural changes within the extensive dopaminergic projections from these regions to the basal ganglia and medial forebrain. Fibers from the SN and striatum also directly innervate the lateral edge of the VA nucleus where we see high isotropy and crossing diffusion orientations (Kultas-ilinsky & Ilinsky, 1990; Sakai et al., 1998). Our findings may signal increased innervation of the thalamus from the SN and/or striatum and/or increased myelination of axons in these pathways. This further suggests that these findings may reflect maturation of cortico-striatal-thalamic pathways that may be important for processes of motor control, cognition and self-regulation that are developing in this age range (Baron Nelson et al., 2019).

Voxels within the GP and putamen showed the largest age associations with N0. These basal ganglia regions form part of parallel frontal, temporal and parietal cortical circuits that are involved in a number of cognitive and motor functions and have a topographical representation across these structures (Alexander et al., 1986; Middleton & Strick, 2000). However, the relatively homogenous developmental effects seen within these regions suggests that these microstructural changes are not associated with any particular topographically organized functional circuit. Despite these similar age related effects across the GP and putamen, overall mean isotropic diffusion across subjects was much larger in the GP which may reflect higher myelin content in the GP compared to the putamen (Lanciego et al., 2012). Throughout adolescence, there is a larger increase in iron concentration, estimated by increased susceptibility on T2* weighted imaging, within the GP relative to the putamen (Larson et al, 2020). This iron related effect has been shown to correlate with DTI metrics in these deep gray matter structures (Pfefferbaum et al., 2010); therefore, increasing iron accumulation may be contributing to the N0 developmental effects that we observe in these regions. However, as developmental changes in N0 were of a similar magnitude across different basal ganglia regions, our results are unlikely to be primarily driven by effects related to iron accumulation. Further research is required to determine the extent to which iron content contributes to these RSI metrics.

Integrated circuit-based theories of adolescent development highlight the importance of both subcortical-subcortical and subcortical-cortical circuitry in the development of multiple important cognitive and behavioral dimensions during this period (Casey et al., 2016). Although we did not measure connectivity in the current study, robust microstructural changes occurring in these subcortical regions may indicate important refinement of these developing circuits. The localization of many of these effects in regions where diffusion primarily occurs in the anterior-posterior direction suggests that there may be differential development along specific pathways (possibly emphasizing fronto-thalamic-striatal circuits) and this may have a direct functional implication for adolescent behavioral development. By using voxelwise analyses of novel RSI measures we have been able to delineate the heterogeneity of microstructural developmental effects across potentially distinct neurite populations. Future work will aim to characterize how these microstructural changes associate with markers of connectivity and the extent to which they can explain individual variability in cognitive and behavioral development.

## ACKNOWLEDGEMENTS

The authors wish to thank the youth and families participating in the Adolescent Brain Cognitive Development (ABCD) Study and all ABCD staff involved in data collection and curation. Dr. Dale reports that he was a Founder of and holds equity in CorTechs Labs, Inc., and serves on its Scientific Advisory Board. He is a member of the Scientific Advisory Board of Human Longevity, Inc. He receives funding through research grants from GE Healthcare to UCSD. The terms of these arrangements have been reviewed by and approved by UCSD in accordance with its conflict of interest policies.

