## Supplementary Material for "Spatially heterogeneous microstructural development within subcortical regions from 9-13 years"

SUPPLEMENTARY TABLES & FIGURES


| | | |
|---|---|---|
| **Supplementary Table 1** | Outline of how different developmental cellular processes can modulate both hindered and restricted diffusion. | Pg 2 |
| **Supplementary Table 2** | Regions of interest (ROIs) labelled using 3 different methods | Pg 3 |
| **Supplementary Table 3** | Summary statistics for the N0 age associations | Pg 4 |
| **Supplementary Table 4** | Summary statistics for the ND age associations | Pg 5 |
| **Supplementary Table 5** | Summary statistics for the MD age associations | Pg 6 |
| **Supplementary Table 6** | Summary statistics for the FA age associations | Pg 7 |
| **Supplementary Figure 1** | Voxelwise statistics for the association between $age_0$ and $age_\Delta$ and restricted diffusion metrics across subcortical regions | Pg 8 |
| **Supplementary Figure 2** | Zoomed in images of the voxelwise $age_\Delta$ thresholded t-statistics and FODs across different brain slices | Pg 9 |
| **Supplementary Figure 3** | Age associations with diffusion tensor metrics across subcortical regions | Pg 10 |
| **Supplementary Figure 4** | Voxelwise statistics for the association between $age_0$ and $age_\Delta$ and diffusion tensor metrics across subcortical regions | Pg 11 |

**Supplementary Table 1. Outline of how different developmental cellular processes can modulate both hindered and restricted diffusion.**

| *Developmental processes* | *Effect on diffusion* | *Hindered signal fraction* | *Restricted signal fraction* |
|---|---|---|---|
| ***Myelination*** | Reduces volume of extracellular space<br>Reduces permeability of axonal membranes, resulting in less exchange of water molecules between intracellular and extracellular spaces | Decrease | Increase |
| ***Increase in neurite diameter with constant neurite density*** | Diameter of neurites will not exceed typical diffusion length scale, therefore will not alter the magnitude of the measured water displacement, but will reduce the volume of extracellular space | Decrease | Increase |
| ***Dendritic sprouting*** | Arborization will reduce the volume of the extracellular space | Decrease | Increase |
| ***Increase in cell size with constant cell density (<<typical diffusion length scale)*** | Reduces volume of extracellular space, therefore increasing the restricted signal fraction, but will not alter the magnitude of the measured water displacement | Decrease | Increase |
| ***Increase in cell size with constant cell density (>>typical diffusion length scale)*** | Restricted signal fraction will increase until >>typical diffusion length scale. Beyond this diffusion will appear hindered | Increase | Decrease |
| ***Increase in number of mature astrocytes, with spongiform morphologies*** | Mature astrocytes have greater permeability relative to neurons causing greater exchange of water molecules between intracellular and extracellular spaces resulting in less restricted diffusion | Increase | Decrease |
| ***Recruitment/Activation of microglia*** | Activated microglia show reduced elongated processes that become thicker, increased cell body size (~8μm), and greater clustering together. This reduces the volume of the extracellular space. | Decrease | Increase |

**Supplementary Table 2. Regions of interest (ROIs) labelled using 3 different methods.** Column 1) automatic segmentation using FreeSurfer 5.3 applied to each subject's T1 image in atlas space (Fischl et al., 2002); Column 2) registration of the Pauli atlas of subcortical nuclei to the multispectral atlas (Pauli et al., 2018); Column 3) registration of the the Najdenovska thalamic nuclei atlas to our data (Najdenovska et al., 2018).

| **FreeSurfer 5.3 segmentation**<br>**T1** | **Pauli, 2018**<br>**HCP T1 & T2** | **Najdenovska, 2018**<br>**HCP FODs** |
|---|---|---|
| Amygdala (Amg) | Putamen (Pu) | Anterior (A) |
| Hippocampus (Hipp) | Caudate (Ca) | Ventral anterior (VA) |
| Putamen (Pu) | Nucleus accumbens (NAcc) | Mediodorsal (tMD) |
| Caudate (Ca) | Extended amygdala (EA) | Ventral-latero-ventral (VLV) |
| Globus pallidus (GP) | Substantia nigra pars compacta (SNpc) | Ventral-latero-dorsal (VLD) |
| Accumbens area (NAcc) | Substantia nigra pars reticulata (SNpr) | Central-latero-lateral-posterior-medial-pulvinar (C) |
| Thalamus (Thal) | Red nucleus (RN) | Pulvinar (P) |
| Ventral Diencephalon (VDC) | Parabrachial pigmented nucleus (PBP) | |
| | Hypothalamus (Hyp) | |
| | Mamillary nucleus (MN) | |
| | Subthalamic nucleus (STN) | |

**Supplementary Table 3. Summary statistics for the N0 age associations.** Statistics include the range of beta coefficients in each ROI for the voxelwise analyses and the estimated beta coefficient, SE, Z statistic and -log10(P) for the ROI analyses. Statistics are shown for $age_0$ and $age_\Delta$ associations and the contrast between these predictors. P-values are adjusted for multiple comparisons (adjusted significance threshold is -log10(P)=1.3).

| | | **$age_0$ voxelwise** | | **$age_0$ ROIwise** | | | | **$age_\Delta$ voxelwise** | | **$age_\Delta$ ROIwise** | | | | **$age_0$ - $age_\Delta$ ROIwise** | | | |
|---|---|---|---|---|---|---|---|---|---|---|---|---|---|---|---|---|---|
| | *ROI* | $min\beta$ | $max\beta$ | $\beta$ | $SE$ | $Z$ | $-\log(P)$ | $min\beta$ | $max\beta$ | $\beta$ | $SE$ | $Z$ | $-\log(P)$ | $\beta$ | $SE$ | $Z$ | $-\log(P)$ |
| ***N0*** | ***Ca*** | -0.002 | 0.01 | 0.02 | 0.001 | 20 | 84 | -0.001 | 0.009 | 0.02 | 0.001 | 36 | 275 | 0 | 0.001 | 0 | -1 |
| ***N0*** | ***Pu*** | 0.003 | 0.012 | 0.03 | 0.001 | 29 | 185 | 0.003 | 0.01 | 0.027 | 0.001 | 44 | Inf | 0.003 | 0.001 | 3 | 1 |
| ***N0*** | ***GP*** | 0.003 | 0.012 | 0.027 | 0.001 | 31 | 203 | 0.003 | 0.011 | 0.025 | 0 | 51 | Inf | 0.003 | 0.001 | 3 | 1 |
| ***N0*** | ***Thal*** | -0.001 | 0.01 | 0.02 | 0.001 | 22 | 107 | -0.001 | 0.01 | 0.02 | 0.001 | 39 | Inf | 0.001 | 0.001 | 1 | -1 |
| ***N0*** | ***NAcc*** | 0.008 | 0.01 | 0.024 | 0.001 | 24 | 124 | 0.007 | 0.009 | 0.022 | 0.001 | 35 | 267 | 0.002 | 0.001 | 2 | 0 |
| ***N0*** | ***VDC*** | -0.002 | 0.012 | 0.021 | 0.001 | 24 | 130 | -0.002 | 0.011 | 0.02 | 0.001 | 38 | 309 | 0.001 | 0.001 | 1 | -1 |
| ***N0*** | ***Hipp*** | 0 | 0.007 | 0.017 | 0.001 | 16 | 57 | 0 | 0.008 | 0.015 | 0.001 | 27 | 155 | 0.002 | 0.001 | 2 | 0 |
| ***N0*** | ***Amy*** | 0.001 | 0.008 | 0.018 | 0.001 | 18 | 70 | -0.001 | 0.008 | 0.015 | 0.001 | 23 | 117 | 0.004 | 0.001 | 3 | 2 |
| ***N0*** | ***SNpc*** | 0.004 | 0.011 | 0.022 | 0.001 | 22 | 107 | 0.004 | 0.01 | 0.021 | 0.001 | 32 | 228 | 0.001 | 0.001 | 1 | -1 |
| ***N0*** | ***SNr*** | 0.006 | 0.011 | 0.024 | 0.001 | 26 | 143 | 0.005 | 0.01 | 0.021 | 0.001 | 37 | 294 | 0.003 | 0.001 | 3 | 1 |
| ***N0*** | ***PBP*** | 0.005 | 0.009 | 0.018 | 0.001 | 17 | 63 | 0.004 | 0.008 | 0.017 | 0.001 | 25 | 141 | 0.001 | 0.001 | 1 | -1 |
| ***N0*** | ***RN*** | 0.002 | 0.008 | 0.015 | 0.001 | 17 | 64 | 0.001 | 0.008 | 0.016 | 0.001 | 29 | 179 | 0 | 0.001 | 0 | -1 |
| ***N0*** | ***Hyp*** | 0.004 | 0.01 | 0.022 | 0.001 | 23 | 114 | 0.002 | 0.01 | 0.021 | 0.001 | 35 | 267 | 0.001 | 0.001 | 1 | -1 |
| ***N0*** | ***MN*** | 0.005 | 0.009 | 0.019 | 0.001 | 18 | 73 | 0.004 | 0.009 | 0.016 | 0.001 | 26 | 146 | 0.003 | 0.001 | 3 | 1 |
| ***N0*** | ***STN*** | 0.001 | 0.008 | 0.015 | 0.001 | 16 | 53 | 0.001 | 0.008 | 0.014 | 0.001 | 23 | 116 | 0.001 | 0.001 | 1 | -1 |
| ***N0*** | ***A*** | 0.003 | 0.009 | 0.018 | 0.001 | 19 | 80 | 0.002 | 0.008 | 0.017 | 0.001 | 33 | 237 | 0.001 | 0.001 | 1 | -1 |
| ***N0*** | ***MD*** | 0.004 | 0.009 | 0.02 | 0.001 | 23 | 111 | 0.003 | 0.009 | 0.02 | 0.001 | 38 | 320 | 0 | 0.001 | 0 | -1 |
| ***N0*** | ***VA*** | 0.003 | 0.01 | 0.022 | 0.001 | 22 | 104 | 0.003 | 0.01 | 0.021 | 0.001 | 38 | 312 | 0.001 | 0.001 | 1 | -1 |
| ***N0*** | ***VLD*** | 0.001 | 0.009 | 0.021 | 0.001 | 21 | 93 | 0.001 | 0.009 | 0.02 | 0.001 | 37 | 291 | 0.001 | 0.001 | 1 | -1 |
| ***N0*** | ***VLV*** | 0.004 | 0.009 | 0.02 | 0.001 | 21 | 96 | 0.002 | 0.009 | 0.019 | 0.001 | 37 | 304 | 0.001 | 0.001 | 1 | -1 |
| ***N0*** | ***CP*** | 0.002 | 0.008 | 0.017 | 0.001 | 18 | 70 | 0.002 | 0.008 | 0.017 | 0.001 | 32 | 223 | -0.001 | 0.001 | -1 | -1 |
| ***N0*** | ***P*** | 0.002 | 0.008 | 0.016 | 0.001 | 16 | 58 | 0.002 | 0.007 | 0.015 | 0.001 | 30 | 201 | 0 | 0.001 | 0 | -1 |

**Supplementary Table 4. Summary statistics for the ND age associations.** Statistics include the range of beta coefficients in each ROI for the voxelwise analyses and the estimated beta coefficient, SE, Z statistic and -log10(P) for the ROI analyses. Statistics are shown for $age_0$ and $age_\Delta$ associations and the contrast between these predictors. P-values are adjusted for multiple comparisons (adjusted significance threshold is -log10(P)=1.3).

| | | $age_0$ voxelwise | | $age_0$ ROIwise | | | | $age_\Delta$ voxelwise | | $age_\Delta$ ROIwise | | | | $age_0$ - $age_\Delta$ ROIwise | | | |
|---|---|---|---|---|---|---|---|---|---|---|---|---|---|---|---|---|---|
| | *ROI* | $min\beta$ | $max\beta$ | $\beta$ | $SE$ | $Z$ | $-\log(P)$ | $min\beta$ | $max\beta$ | $\beta$ | $SE$ | $Z$ | $-\log(P)$ | $\beta$ | $SE$ | $Z$ | $-\log(P)$ |
| ***ND*** | ***Ca*** | -0.001 | 0.006 | 0.007 | 0.001 | 9 | 17 | -0.002 | 0.007 | 0.007 | 0 | 18 | 67 | 0 | 0.001 | 0 | -1 |
| ***ND*** | ***Pu*** | -0.001 | 0.006 | 0.013 | 0.001 | 13 | 38 | -0.002 | 0.006 | 0.012 | 0 | 28 | 175 | 0.001 | 0.001 | 1 | -1 |
| ***ND*** | ***GP*** | -0.001 | 0.006 | 0.011 | 0.001 | 12 | 29 | -0.001 | 0.006 | 0.012 | 0.001 | 23 | 117 | -0.001 | 0.001 | -1 | -1 |
| ***ND*** | ***Thal*** | -0.002 | 0.008 | 0.016 | 0.001 | 14 | 43 | -0.001 | 0.007 | 0.018 | 0.001 | 35 | 273 | -0.002 | 0.001 | -2 | 0 |
| ***ND*** | ***NAcc*** | 0.002 | 0.006 | 0.013 | 0.001 | 13 | 38 | 0.001 | 0.006 | 0.012 | 0.001 | 24 | 124 | 0.001 | 0.001 | 1 | -1 |
| ***ND*** | ***VDC*** | -0.001 | 0.009 | 0.019 | 0.001 | 18 | 72 | -0.001 | 0.009 | 0.022 | 0.001 | 41 | Inf | -0.003 | 0.001 | -3 | 1 |
| ***ND*** | ***Hipp*** | -0.002 | 0.005 | 0.008 | 0.001 | 8 | 13 | -0.002 | 0.006 | 0.007 | 0 | 16 | 55 | 0.001 | 0.001 | 1 | -1 |
| ***ND*** | ***Amy*** | -0.002 | 0.006 | 0.007 | 0.001 | 7 | 10 | -0.002 | 0.006 | 0.007 | 0 | 16 | 52 | 0 | 0.001 | 0 | -1 |
| ***ND*** | ***SNpc*** | 0.003 | 0.008 | 0.02 | 0.001 | 17 | 59 | 0.003 | 0.009 | 0.021 | 0.001 | 36 | 287 | -0.001 | 0.001 | -1 | -1 |
| ***ND*** | ***SNr*** | 0.001 | 0.008 | 0.015 | 0.001 | 14 | 40 | 0.002 | 0.008 | 0.016 | 0.001 | 29 | 187 | -0.001 | 0.001 | -1 | -1 |
| ***ND*** | ***PBP*** | 0.002 | 0.006 | 0.016 | 0.001 | 15 | 47 | 0.003 | 0.007 | 0.02 | 0.001 | 29 | 180 | -0.003 | 0.001 | -3 | 1 |
| ***ND*** | ***RN*** | 0.001 | 0.005 | 0.012 | 0.001 | 10 | 22 | 0.001 | 0.006 | 0.016 | 0.001 | 24 | 121 | -0.004 | 0.001 | -3 | 1 |
| ***ND*** | ***Hyp*** | -0.001 | 0.008 | 0.016 | 0.001 | 14 | 44 | -0.001 | 0.009 | 0.019 | 0.001 | 30 | 202 | -0.002 | 0.001 | -2 | 0 |
| ***ND*** | ***MN*** | 0.003 | 0.008 | 0.016 | 0.001 | 13 | 37 | 0.003 | 0.008 | 0.018 | 0.001 | 28 | 170 | -0.001 | 0.001 | -1 | -1 |
| ***ND*** | ***STN*** | 0 | 0.003 | 0.005 | 0.001 | 6 | 6 | 0 | 0.003 | 0.007 | 0.001 | 12 | 30 | -0.002 | 0.001 | -2 | 0 |
| ***ND*** | ***A*** | 0 | 0.007 | 0.013 | 0.001 | 11 | 27 | 0 | 0.007 | 0.014 | 0.001 | 25 | 132 | -0.001 | 0.001 | -1 | -1 |
| ***ND*** | ***MD*** | 0.001 | 0.007 | 0.016 | 0.001 | 13 | 39 | 0.001 | 0.007 | 0.017 | 0.001 | 29 | 178 | -0.001 | 0.001 | -1 | -1 |
| ***ND*** | ***VA*** | 0.001 | 0.008 | 0.018 | 0.001 | 16 | 55 | 0.001 | 0.007 | 0.018 | 0.001 | 34 | 255 | 0 | 0.001 | 0 | -1 |
| ***ND*** | ***VLD*** | -0.001 | 0.006 | 0.012 | 0.001 | 10 | 21 | 0.001 | 0.007 | 0.016 | 0.001 | 29 | 187 | -0.003 | 0.001 | -2 | 0 |
| ***ND*** | ***VLV*** | 0 | 0.007 | 0.007 | 0.001 | 6 | 6 | 0 | 0.006 | 0.01 | 0.001 | 15 | 46 | -0.002 | 0.001 | -2 | 0 |
| ***ND*** | ***CP*** | 0 | 0.006 | 0.007 | 0.001 | 6 | 8 | 0 | 0.007 | 0.009 | 0 | 20 | 89 | -0.002 | 0.001 | -1 | -1 |
| ***ND*** | ***P*** | -0.002 | 0.005 | 0.012 | 0.001 | 9 | 18 | -0.002 | 0.005 | 0.013 | 0.001 | 24 | 127 | -0.001 | 0.001 | -1 | -1 |

**Supplementary Table 5. Summary statistics for the MD age associations.** Statistics include the range of beta coefficients in each ROI for the voxelwise analyses and the estimated beta coefficient, SE, Z statistic and -log10(P) for the ROI analyses. Statistics are shown for $age_0$ and $age_\Delta$ associations and the contrast between these predictors. P-values are adjusted for multiple comparisons (adjusted significance threshold is -log10(P)=1.3).

| | | **$age_0$ voxelwise** | | **$age_0$ ROIwise** | | | | **$age_\Delta$ voxelwise** | | **$age_\Delta$ ROIwise** | | | | **$age_0$ - $age_\Delta$ ROIwise** | | | |
|---|---|---|---|---|---|---|---|---|---|---|---|---|---|---|---|---|---|
| | *ROI* | $min\beta$ | $max\beta$ | $\beta$ | $SE$ | $Z$ | $-\log(P)$ | $min\beta$ | $max\beta$ | $\beta$ | $SE$ | $Z$ | $-\log(P)$ | $\beta$ | $SE$ | $Z$ | $-\log(P)$ |
| ***MD*** | ***Ca*** | -0.01 | 0.001 | -0.017 | 0.001 | -20 | 89 | -0.01 | 0.001 | -0.017 | 0 | -37 | 291 | 0 | 0.001 | 0 | -1 |
| ***MD*** | ***Pu*** | -0.012 | -0.003 | -0.025 | 0.001 | -30 | 189 | -0.012 | -0.004 | -0.025 | 0 | -53 | Inf | 0 | 0.001 | 0 | -1 |
| ***MD*** | ***GP*** | -0.011 | -0.005 | -0.024 | 0.001 | -31 | 209 | -0.01 | -0.005 | -0.023 | 0 | -52 | Inf | -0.001 | 0.001 | -2 | 0 |
| ***MD*** | ***Thal*** | -0.009 | 0 | -0.017 | 0.001 | -24 | 123 | -0.01 | 0 | -0.018 | 0 | -48 | Inf | 0.001 | 0.001 | 1 | -1 |
| ***MD*** | ***NAcc*** | -0.009 | -0.006 | -0.021 | 0.001 | -25 | 133 | -0.008 | -0.006 | -0.019 | 0.001 | -36 | 280 | -0.002 | 0.001 | -2 | 0 |
| ***MD*** | ***VDC*** | -0.01 | -0.002 | -0.019 | 0.001 | -28 | 169 | -0.011 | -0.001 | -0.02 | 0 | -50 | Inf | 0.001 | 0.001 | 2 | 0 |
| ***MD*** | ***Hipp*** | -0.007 | -0.001 | -0.014 | 0.001 | -17 | 62 | -0.008 | 0 | -0.013 | 0 | -29 | 182 | -0.001 | 0.001 | -2 | 0 |
| ***MD*** | ***Amy*** | -0.008 | -0.003 | -0.017 | 0.001 | -19 | 76 | -0.01 | -0.001 | -0.015 | 0.001 | -29 | 177 | -0.002 | 0.001 | -2 | 0 |
| ***MD*** | ***SNpc*** | -0.009 | -0.005 | -0.02 | 0.001 | -26 | 148 | -0.01 | -0.005 | -0.021 | 0 | -42 | Inf | 0 | 0.001 | 0 | -1 |
| ***MD*** | ***SNr*** | -0.009 | -0.006 | -0.021 | 0.001 | -27 | 157 | -0.009 | -0.006 | -0.02 | 0 | -45 | Inf | 0 | 0.001 | 0 | -1 |
| ***MD*** | ***PBP*** | -0.008 | -0.006 | -0.018 | 0.001 | -21 | 93 | -0.008 | -0.007 | -0.018 | 0.001 | -33 | 239 | 0 | 0.001 | 0 | -1 |
| ***MD*** | ***RN*** | -0.009 | -0.005 | -0.018 | 0.001 | -21 | 96 | -0.011 | -0.005 | -0.021 | 0.001 | -41 | Inf | 0.004 | 0.001 | 4 | 2 |
| ***MD*** | ***Hyp*** | -0.01 | -0.003 | -0.019 | 0.001 | -24 | 121 | -0.011 | -0.002 | -0.018 | 0 | -37 | 296 | -0.001 | 0.001 | -1 | -1 |
| ***MD*** | ***MN*** | -0.008 | -0.005 | -0.017 | 0.001 | -20 | 89 | -0.008 | -0.003 | -0.014 | 0 | -30 | 189 | -0.002 | 0.001 | -3 | 1 |
| ***MD*** | ***STN*** | -0.009 | -0.004 | -0.016 | 0.001 | -19 | 78 | -0.009 | -0.004 | -0.017 | 0.001 | -32 | 216 | 0.001 | 0.001 | 1 | -1 |
| ***MD*** | ***A*** | -0.008 | -0.003 | -0.015 | 0.001 | -20 | 85 | -0.008 | -0.002 | -0.016 | 0 | -36 | 289 | 0 | 0.001 | 0 | -1 |
| ***MD*** | ***MD*** | -0.008 | -0.003 | -0.017 | 0.001 | -25 | 131 | -0.009 | -0.003 | -0.018 | 0 | -47 | Inf | 0.001 | 0.001 | 2 | 0 |
| ***MD*** | ***VA*** | -0.008 | -0.003 | -0.018 | 0.001 | -22 | 109 | -0.009 | -0.003 | -0.019 | 0 | -45 | Inf | 0 | 0.001 | 0 | -1 |
| ***MD*** | ***VLD*** | -0.008 | -0.001 | -0.017 | 0.001 | -21 | 100 | -0.009 | 0 | -0.019 | 0 | -49 | Inf | 0.001 | 0.001 | 2 | 0 |
| ***MD*** | ***VLV*** | -0.009 | -0.005 | -0.018 | 0.001 | -22 | 104 | -0.01 | -0.004 | -0.02 | 0 | -45 | Inf | 0.001 | 0.001 | 1 | 0 |
| ***MD*** | ***CP*** | -0.007 | -0.002 | -0.015 | 0.001 | -20 | 86 | -0.008 | -0.002 | -0.016 | 0 | -40 | Inf | 0.002 | 0.001 | 2 | 0 |
| ***MD*** | ***P*** | -0.007 | -0.003 | -0.014 | 0.001 | -18 | 70 | -0.008 | -0.002 | -0.015 | 0 | -37 | 305 | 0.001 | 0.001 | 1 | -1 |

**Supplementary Table 6. Summary statistics for the FA age associations.** Statistics include the range of beta coefficients in each ROI for the voxelwise analyses and the estimated beta coefficient, SE, Z statistic and -log10(P) for the ROI analyses. Statistics are shown for $age_0$ and $age_\Delta$ associations and the contrast between these predictors. P-values are adjusted for multiple comparisons (adjusted significance threshold is -log10(P)=1.3).

| | | **$age_0$ voxelwise** | | **$age_0$ ROIwise** | | | | **$age_\Delta$ voxelwise** | | **$age_\Delta$ ROIwise** | | | | **$age_0$ - $age_\Delta$ ROIwise** | | | |
|---|---|---|---|---|---|---|---|---|---|---|---|---|---|---|---|---|---|
| | *ROI* | $min\beta$ | $max\beta$ | $\beta$ | $SE$ | $Z$ | $-\log(P)$ | $min\beta$ | $max\beta$ | $\beta$ | $SE$ | $Z$ | $-\log(P)$ | $\beta$ | $SE$ | $Z$ | $-\log(P)$ |
| ***FA*** | ***Ca*** | -0.002 | 0.005 | 0.005 | 0.001 | 5 | 5 | -0.001 | 0.006 | 0.008 | 0 | 19 | 76 | -0.003 | 0.001 | -3 | 1 |
| ***FA*** | ***Pu*** | -0.003 | 0.005 | 0.006 | 0.001 | 7 | 9 | -0.003 | 0.004 | 0.009 | 0 | 21 | 99 | -0.002 | 0.001 | -2 | 0 |
| ***FA*** | ***GP*** | -0.003 | 0.003 | 0 | 0.001 | 0 | -1 | -0.003 | 0.005 | 0.003 | 0 | 6 | 7 | -0.003 | 0.001 | -3 | 1 |
| ***FA*** | ***Thal*** | -0.002 | 0.005 | 0.007 | 0.001 | 8 | 14 | -0.001 | 0.005 | 0.011 | 0 | 27 | 159 | -0.004 | 0.001 | -4 | 3 |
| ***FA*** | ***NAcc*** | -0.001 | 0.004 | 0.006 | 0.001 | 6 | 8 | 0 | 0.004 | 0.007 | 0.001 | 13 | 37 | -0.001 | 0.001 | -1 | -1 |
| ***FA*** | ***VDC*** | -0.004 | 0.006 | 0.011 | 0.001 | 13 | 38 | -0.003 | 0.007 | 0.016 | 0 | 36 | 288 | -0.005 | 0.001 | -5 | 5 |
| ***FA*** | ***Hipp*** | -0.003 | 0.005 | 0.004 | 0.001 | 4 | 3 | -0.003 | 0.005 | 0.005 | 0 | 11 | 25 | -0.001 | 0.001 | -1 | -1 |
| ***FA*** | ***Amy*** | -0.003 | 0.004 | 0.002 | 0.001 | 2 | 0 | -0.003 | 0.005 | 0.005 | 0 | 11 | 24 | -0.003 | 0.001 | -2 | 0 |
| ***FA*** | ***SNpc*** | 0.002 | 0.005 | 0.013 | 0.001 | 13 | 36 | 0.003 | 0.005 | 0.016 | 0.001 | 31 | 206 | -0.003 | 0.001 | -3 | 1 |
| ***FA*** | ***SNr*** | 0 | 0.005 | 0.009 | 0.001 | 9 | 17 | 0.001 | 0.005 | 0.012 | 0.001 | 23 | 114 | -0.003 | 0.001 | -3 | 1 |
| ***FA*** | ***PBP*** | 0.001 | 0.004 | 0.012 | 0.001 | 12 | 31 | 0.002 | 0.005 | 0.016 | 0.001 | 25 | 141 | -0.004 | 0.001 | -4 | 3 |
| ***FA*** | ***RN*** | 0 | 0.005 | 0.01 | 0.001 | 10 | 20 | 0.001 | 0.005 | 0.014 | 0.001 | 24 | 122 | -0.004 | 0.001 | -3 | 2 |
| ***FA*** | ***Hyp*** | -0.004 | 0.006 | 0.007 | 0.001 | 8 | 14 | -0.003 | 0.007 | 0.011 | 0 | 23 | 117 | -0.004 | 0.001 | -4 | 3 |
| ***FA*** | ***MN*** | 0.001 | 0.005 | 0.01 | 0.001 | 10 | 20 | 0.002 | 0.006 | 0.013 | 0.001 | 21 | 95 | -0.003 | 0.001 | -2 | 1 |
| ***FA*** | ***STN*** | -0.001 | 0.003 | 0.005 | 0.001 | 5 | 4 | 0 | 0.003 | 0.007 | 0.001 | 11 | 26 | -0.002 | 0.001 | -2 | 0 |
| ***FA*** | ***A*** | -0.001 | 0.005 | 0.006 | 0.001 | 6 | 8 | 0 | 0.005 | 0.008 | 0 | 17 | 59 | -0.002 | 0.001 | -2 | 0 |
| ***FA*** | ***MD*** | -0.001 | 0.004 | 0.007 | 0.001 | 8 | 13 | 0 | 0.005 | 0.01 | 0 | 21 | 99 | -0.003 | 0.001 | -3 | 1 |
| ***FA*** | ***VA*** | 0 | 0.005 | 0.01 | 0.001 | 11 | 26 | 0.001 | 0.005 | 0.012 | 0 | 26 | 149 | -0.002 | 0.001 | -2 | 0 |
| ***FA*** | ***VLD*** | -0.001 | 0.004 | 0.006 | 0.001 | 6 | 8 | 0 | 0.005 | 0.011 | 0 | 23 | 117 | -0.005 | 0.001 | -4 | 4 |
| ***FA*** | ***VLV*** | -0.001 | 0.004 | 0.001 | 0.001 | 1 | -1 | -0.001 | 0.004 | 0.004 | 0.001 | 8 | 13 | -0.003 | 0.001 | -3 | 1 |
| ***FA*** | ***CP*** | -0.002 | 0.004 | 0.001 | 0.001 | 1 | -1 | -0.001 | 0.004 | 0.004 | 0 | 10 | 21 | -0.003 | 0.001 | -3 | 1 |
| ***FA*** | ***P*** | -0.003 | 0.003 | 0.005 | 0.001 | 5 | 5 | -0.003 | 0.004 | 0.007 | 0 | 16 | 53 | -0.002 | 0.001 | -2 | 0 |

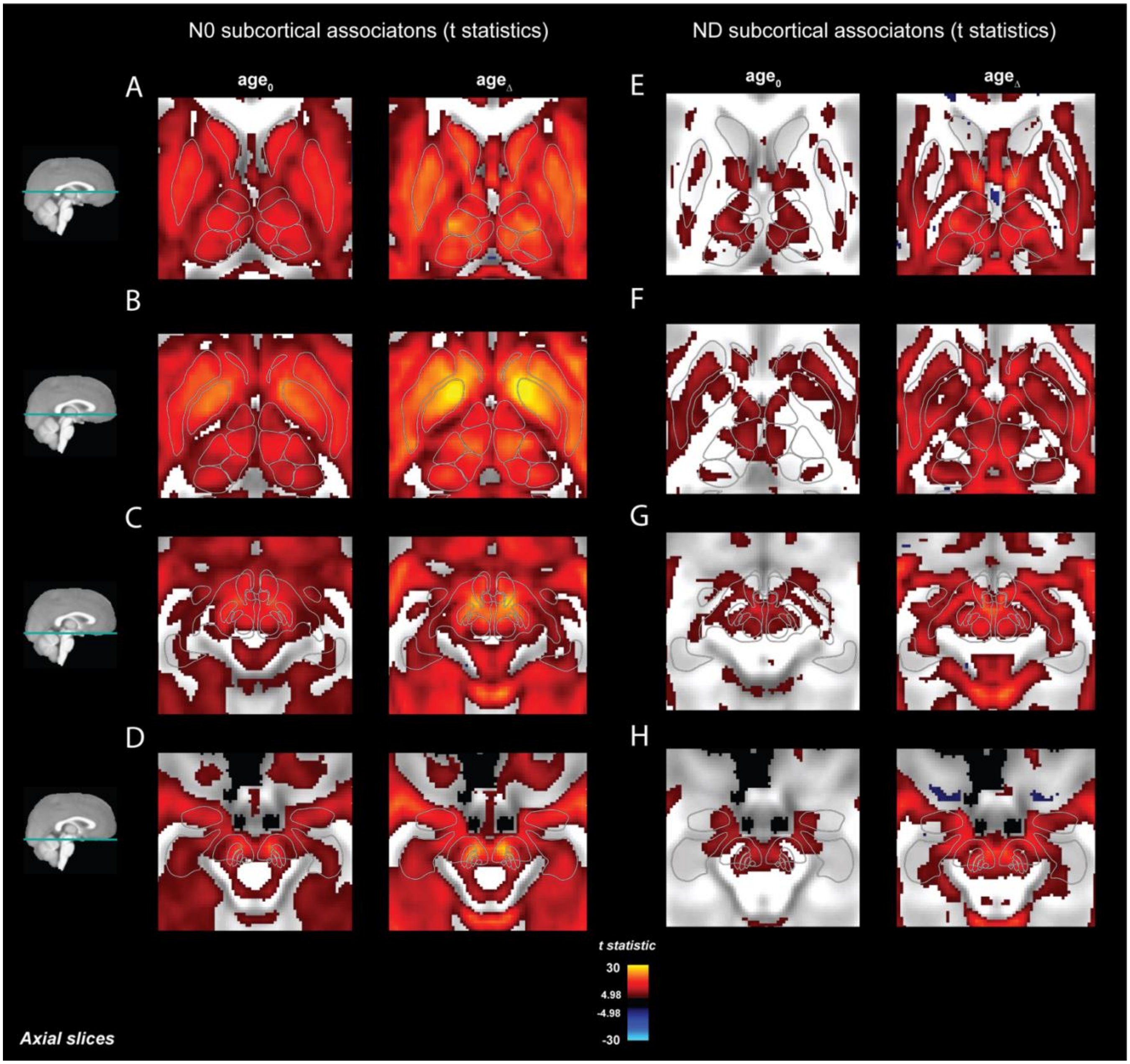


**Supplementary Figure 1. Voxelwise statistics for the association between $age_0$ and $age_\Delta$ and restricted diffusion metrics across subcortical regions.** A-D) T-statistics thresholded at t>|4.98| for the association between N0 and $age_0$ (left) and $age_\Delta$ (right). E-H) T-statistics thresholded at t>|4.98| for the association between ND and $age_0$ (left) and $age_\Delta$ (right). The threshold was determined using a Bonferroni correction for multiple comparisons at an alpha level of 0.05 across all voxels. Outlines of the Aseg, Pauli and Najdenovska ROIs are overlaid. In general, t-statistics were larger for longitudinal associations.

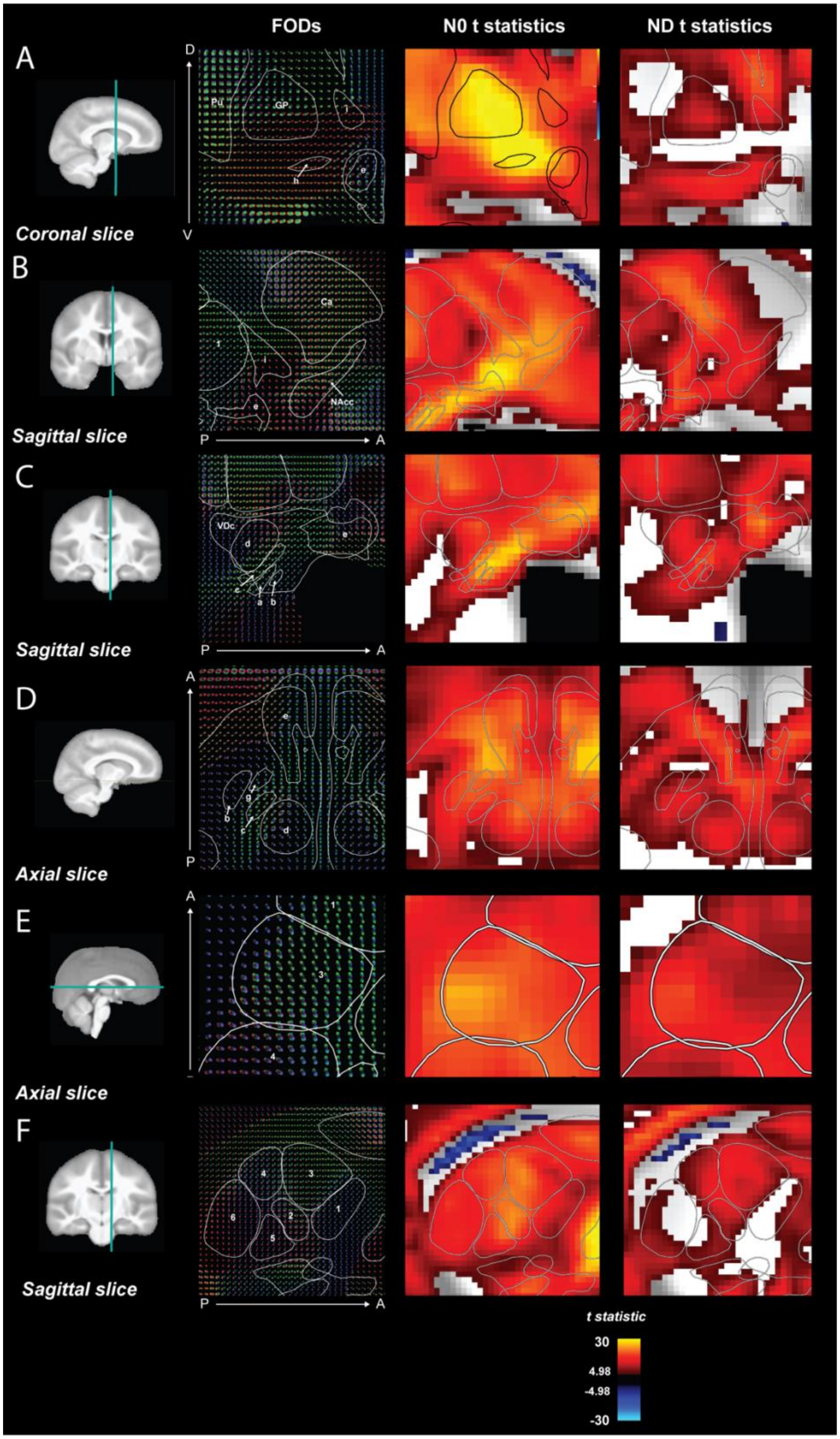


**Supplementary Figure 2. Zoomed in images of the voxelwise age$_\Delta$ thresholded t-statistics and FODs across different brain slices**. T-statistics thresholded at t>|4.98|. The threshold was determined using a Bonferroni correction for multiple comparisons at an alpha level of 0.05 across all voxels. See Figure 3 legend for details of each panal.

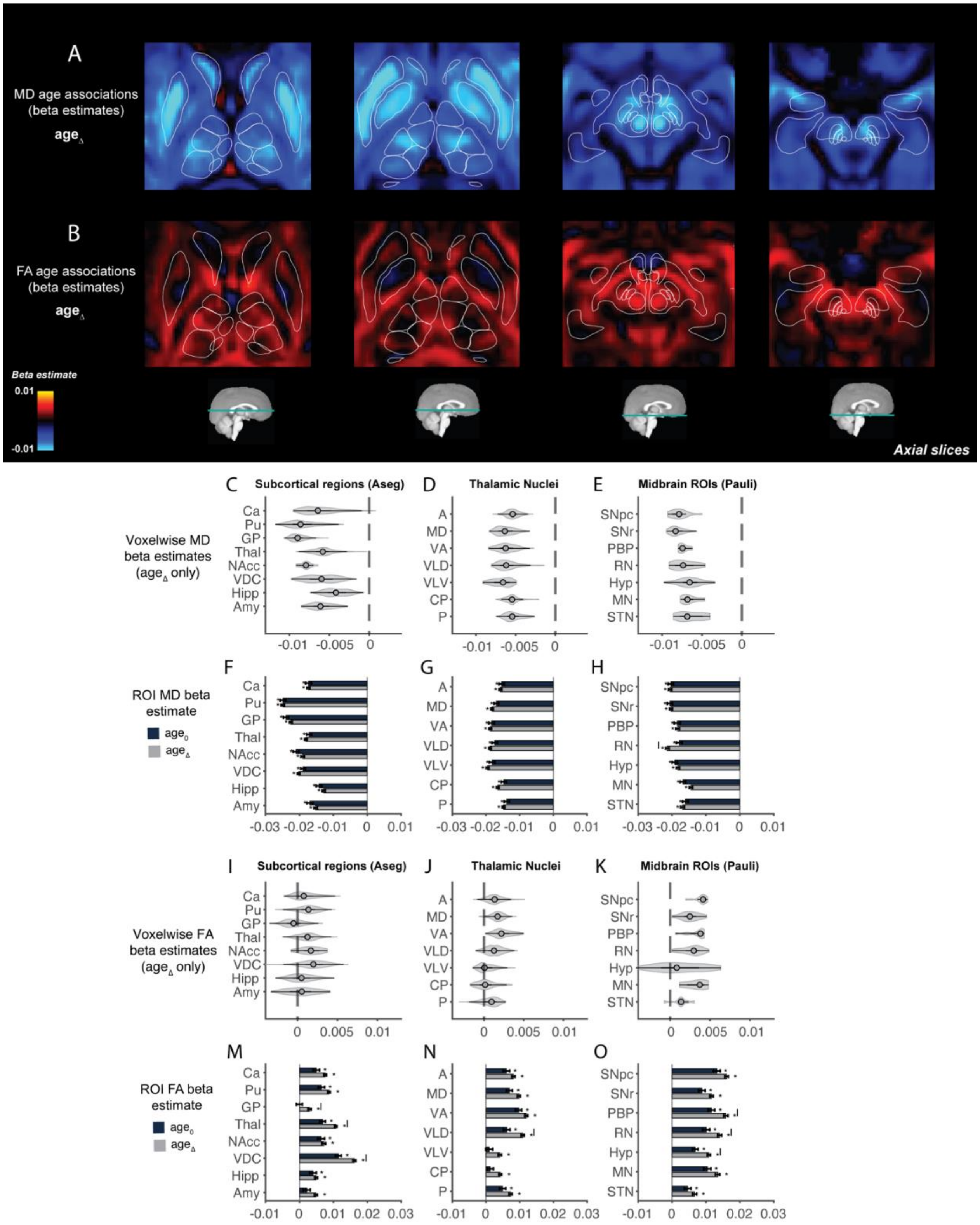


**Supplementary Figure 3. Age associations with diffusion tensor metrics across subcortical regions.** Voxelwise beta coefficients for the age$_\Delta$ associations across different axial brain slices moving from superior (left) to inferior (right) with MD (A) and FA (B). Outlines of the Aseg, Pauli and Najdenovska ROIs are overlaid. Violin plots showing the distribution of voxelwise age$_\Delta$ associations in each ROI for MD (C-E) and FA (I-K) and bar plots of ROI analyses for age$_0$ (dark blue) and age$_\Delta$ (gray) for MD (F-H) and FA (M-O).

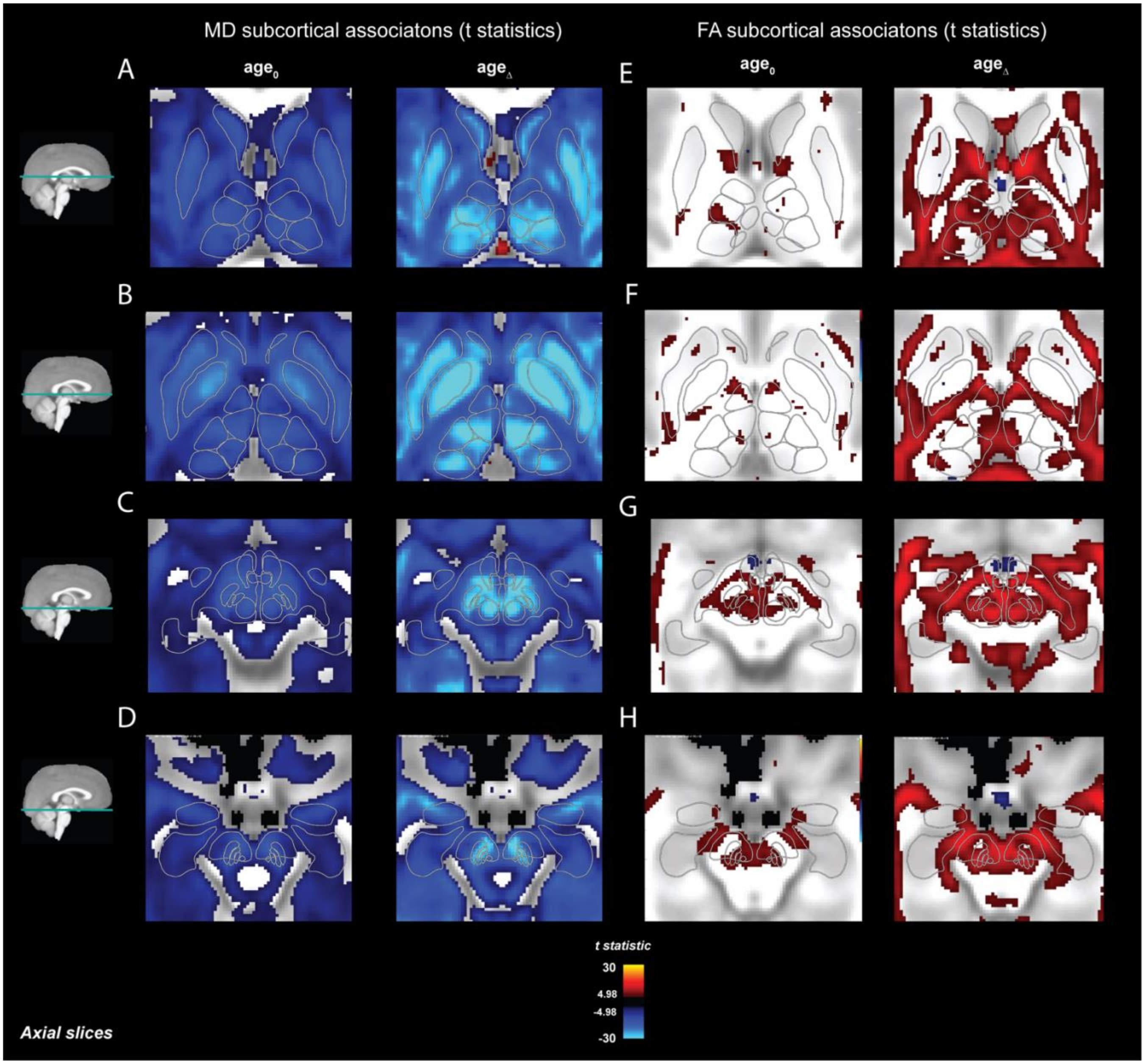


**Supplementary Figure 4. Voxelwise statistics for the association between $age_0$ and $age_\Delta$ and diffusion tensor metrics across subcortical regions.** A-D) T-statistics thresholded at t>|4.98| for the association between MD and $age_0$ (left) and $age_\Delta$ (right). E-H) T-statistics thresholded at t>|4.98| for the association between FA and $age_0$ (left) and $age_\Delta$ (right). The threshold was determined using a Bonferroni correction for multiple comparisons at an alpha level of 0.05 across all voxels. Outlines of the Aseg, Pauli and Najdenovska ROIs are overlaid. In general, t-statistics were larger for longitudinal associations.
